# Lhcf9 is a novel negative regulator of non-photochemical quenching in the diatom *Chaetoceros gracilis*

**DOI:** 10.64898/2026.02.06.704175

**Authors:** Midori Nakamura, Minoru Kumazawa, Ryo Nagao, Takehiro Suzuki, Shoko Tsuji, Hazuki Hasegawa, Hiroaki Takebe, Atsushi Sakurai, Sousuke Imamura, Noriko Ishikawa, Naoshi Dohmae, Seiji Akimoto, Kentaro Ifuku

## Abstract

Photosynthetic organisms in aquatic environments experience rapid fluctuations in light intensity, light quality, and carbon availability, requiring tight regulation of photosynthetic energy conversion. In marine diatoms, non-photochemical quenching (NPQ), particularly energy-dependent quenching (qE), plays a central role in dissipating excess excitation energy as heat. However, excessive NPQ can reduce photosynthetic efficiency under light-limiting or carbon-rich conditions, and how this trade-off is regulated remains poorly understood. Here, we identify CgLhcf9, a previously uncharacterized light-harvesting complex (LHC) protein, as a negative regulator of qE-type NPQ in the centric diatom *Chaetoceros gracilis*. Expression of CgLhcf9 is induced under low red-light and high CO_2_ conditions and strongly suppressed by blue light, indicating regulation by both light quality and carbon availability. Functional analyses using CgLhcf9 knockout and overexpression lines reveal that CgLhcf9 suppresses qE: NPQ induction is enhanced in the absence of CgLhcf9, whereas its accumulation downregulates NPQ without affecting other established qE effectors, including Lhcx1 or xanthophyll cycle pigments. Notably, CgLhcf9 accumulation improves cellular growth under light-limiting conditions. These results identify CgLhcf9 as a novel LHC-type regulator that fine-tunes photosynthetic energy dissipation in response to environmental signals. Our findings establish a regulatory mechanism that balances photoprotection, electron transport, and carbon fixation, advancing our understanding of how marine diatoms optimize photosynthesis under fluctuating light and CO_2_ conditions.

**Significance statement:** Photosynthetic microalgae must balance light-driven electron transport with carbon fixation to maximize growth under fluctuating light and CO_2_ conditions. While non-photochemical quenching (NPQ) protects photosystems from excess light, excessive NPQ can limit photosynthetic efficiency when light or carbon is limiting. Here, we identify the antenna protein CgLhcf9 as a negative regulator of energy-dependent NPQ in the marine diatom *Chaetoceros gracilis*. CgLhcf9 integrates light-quality and CO_2_ signals to suppress NPQ without altering canonical quenching effectors, thereby improving growth under light-limiting conditions. This study reveals a regulatory role for a light-harvesting complex protein in tuning the balance between photoprotection and photosynthetic efficiency, providing insight into how marine diatoms coordinate electron transport and carbon fixation in dynamic environments.

## Introduction

In aquatic environments, both the intensity and spectral quality of light change with depth due to absorption and scattering. In pure water, red light attenuates rapidly, while blue and green wavelengths penetrate deeper (1). In natural seawater, however, dissolved substances and suspended matter further modify light transmission; as their concentration increases, the proportion of blue light generally declines, while green and red wavelengths become more prominent (2). Additionally, phytoplankton community composition can further alter the underwater light spectrum (3). Beyond these depth-dependent changes, vertical mixing causes cells to move through the water column, exposing them to rapid fluctuations in light intensity. Weather conditions also affect surface irradiance. Adapting to such dynamic light environments is a critical survival strategy for photosynthetic organisms, and the capacity to effectively utilize the limited spectrum of underwater light is essential to their ecological success.

Diatoms are one of the successful algae that are responsible for approximately 20% of global carbon fixation, highlighting their role as major primary producers in aquatic ecosystems (4). In addition to carbon, diatoms also play central roles in the biogeochemical cycling of nutrients such as nitrogen and silicon (5). One of the key traits supporting the ecological success of diatoms is the remarkable flexibility of their photosynthetic machinery to adjust to varying underwater light conditions (6). Adaptation to the light environment requires precise regulation of the molecular machinery surrounding the photosynthetic apparatus, with Light-Harvesting Complex (LHC) proteins playing a central role in this process. LHCs are classified into multiple functional subtypes of pigment-protein complexes, some of which function as antenna molecules that efficiently capture light and transfer the excitation energy to either photosystem I (PSI) or photosystem II (PSII). In contrast, LHCs are also involved in dissipating excess light energy as heat via non-photochemical quenching (NPQ), particularly energy-dependent quenching (qE), which serves to protect the photosynthetic machinery from photodamage (7–9).

In green plants, qE-NPQ involves the concerted action of LHC proteins, the xanthophyll cycle, and PsbS and/or Lhcsr protein (10, 11). In diatoms, a similar yet distinct system operates, in which qE is mediated by Lhcx proteins together with their specific xanthophyll cycle, involving the reversible conversion between diadinoxanthin (Ddx) and diatoxanthin (Dtx), responding sensitively to changes in light intensity (12–14). The qE plays a crucial role in maintaining photosynthetic efficiency under fluctuating light conditions by preventing the overexcitation of photosystems and minimizing the generation of reactive oxygen species (ROS). On the other hand, qE in excess can downregulate photosynthesis under light-limiting conditions (15). Therefore, the fine-tuned regulation of the qE component is essential to maintain optimal photosynthesis, especially in dynamically changing light environments where sudden irradiance shifts frequently occur. To achieve this, photosynthetic organisms need to regulate the accumulation and activity of the qE effectors in response to changes in the light environment (16).

In diatoms, the LHCs are phylogenetically distinct from those found in green lineage organisms and bind chlorophyll (chl) *c* and the carotenoid fucoxanthin in addition to chl *a* (7). Fucoxanthin and chl *c* absorb strongly in the blue and green spectral regions, making them well-suited for efficient light capture in aquatic environments (17). Due to this feature, LHCs in diatoms are referred to as fucoxanthin chl *a*/*c*-binding proteins (FCPs). *Chaetoceros* is one of the most abundant genera of marine centric diatoms, and *C. gracilis* is particularly advantageous for the functional analysis of LHCs in diatoms, because of its publicly available genome sequence (ChaetoBase: https://chaetoceros.nibb.ac.jp/) and previously resolved photosystem complex structures (18–20). Genomic and transcriptomic analyses of *C. gracilis* have identified 46 distinct LHCs, which are classified into six subfamilies: Lhcf, Lhcr, Lhcz, Lhcx, Lhcq, and a group of CgLhcr9 homologs (21). Among these, Lhcr and the newly identified Lhcq subfamily are mainly associated with PSI, while Lhcf proteins are primarily bound to PSII. The Lhcf subfamily exhibits notable diversification among species, suggesting its potential role in facilitating environmental adaptation. In addition, there are three *Lhcx* genes in *C. gracilis,* and CgLhcx1 is a major isoform regulating NPQ (14).

Interestingly, when *C. gracilis* is cultured under red light, a novel band of fucoxanthin chl *a*/*c*-binding protein (FCP) has been reported to appear on clear native polyacrylamide gel electrophoresis (CN-PAGE) (22). This band is also observed under specific conditions with elevated CO_2_ levels (23). As for red-light-induced FCPs in diatoms, PtLhcf15 has been characterized in a model pennate diatom *Phaeodactylum tricornutum* (3, 24). PtLhcf15 is suggested to be involved in complementary chromatic adaptation, while its regulation by CO_2_ levels has not been reported. In this study, the red-light and high CO_2_ inducible LHC in *C. gracilis* was identified as CgLhcf9 and has been revealed to be a novel LHC that negatively regulates the qE-NPQ in this diatom.

## Results

### Identification of a Distinct FCP (CgLhcf9) Induced Under Specific Environmental Conditions

To identify a unique FCP that accumulates specifically under red light and high CO_2_ conditions, thylakoid membranes were isolated from cells cultured under each condition and subjected to CN-PAGE. Figures 1A and 1D show the specific bands observed in thylakoid membranes isolated from the cells cultured under red-light illumination with normal air bubbling (RL/Air) or under white-light illumination with 3% CO_2_ bubbling at 30 °C (WL/CO_2_/30 °C). These bands were previously designated as the F2 band (22). The gel strips of CN-PAGE were then excised and subsequently analyzed by SDS-PAGE in the second dimension (Fig. 1B and 1E). The resulting gel showed a protein spot corresponding to the previously designated as FCP-D region (22). Analysis using tandem mass spectrometry suggests that the F2 band—in both red light and elevated CO_2_ conditions—contains a single LHC isoform, CgLhcf9 (Fig. 1C and 1F). CgLhcf9 belongs to a distinct clade in the phylogenetic tree of the light-harvesting complex (LHC) proteins in diatoms (21). In this clade, PtLhcf15 has been known for its red-light inducibility and emission of long-wavelength fluorescence at 710 nm (F710) (3, 24). In addition, PtLhcf13 and PtLhcf14 in *P. tricornutum* are also members of this group, although their gene expression and function have not been well understood.

**Figure 1.**
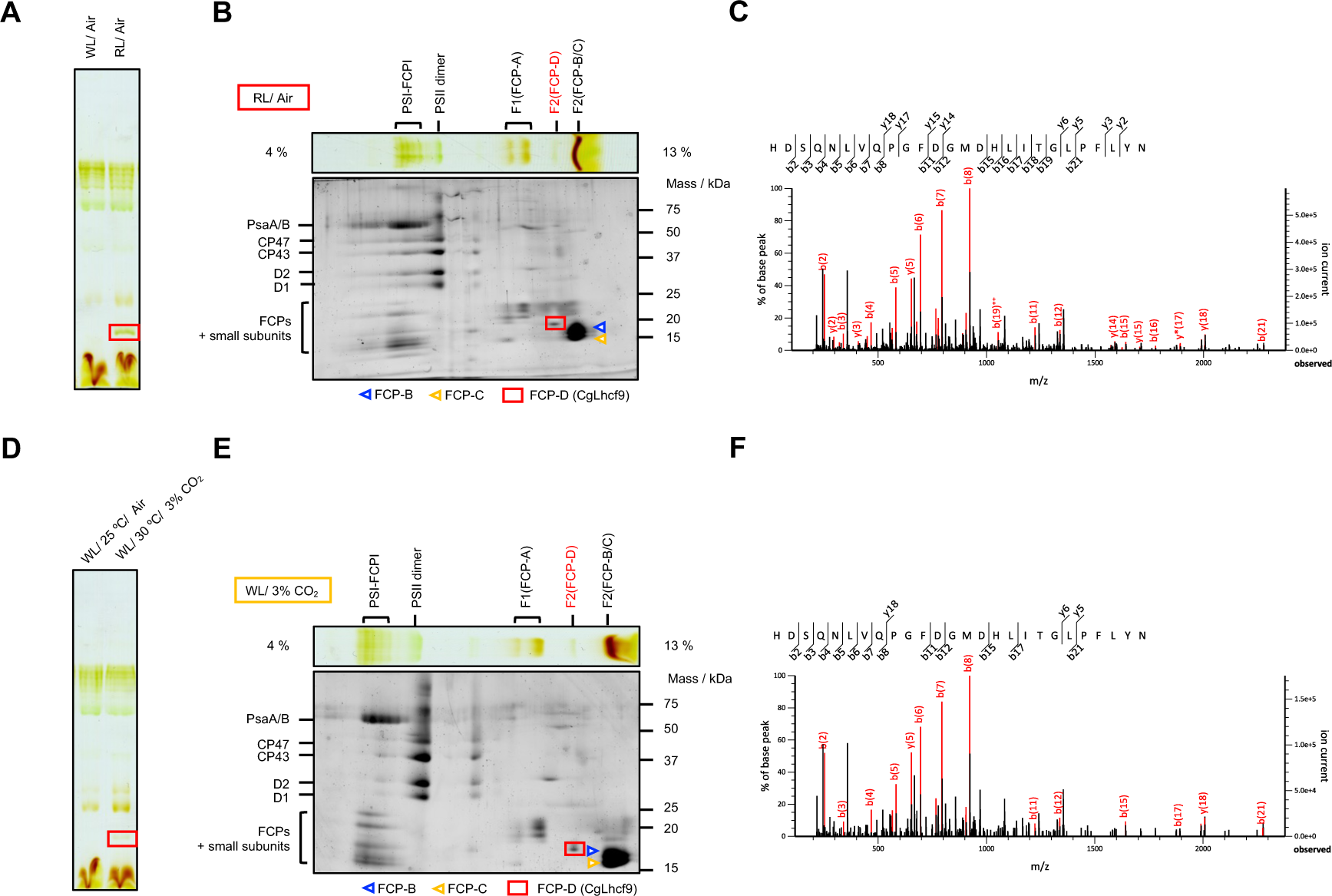
Identification of FCP responding to light and CO_2_ conditions in Chaetoceros gracilis. (A) Comparison of thylakoid membranes isolated under 25 °C, Air conditions with white light and red-light treatments by CN-PAGE. Thylakoid membranes were solubilized with 2% β-DDM, and the cathode buffer was supplemented with 2% Triton X-100 and 2% DOC. (B) Results of two-dimensional SDS-PAGE performed following CN-PAGE of thylakoid membranes obtained under 25 °C, Air, red-light conditions. (C)The FCP spot from the F2 complex was analyzed by mass spectrometry. A representative MS/MS spectrum of the peptides of Lhcf9 in the 25/air/R cells. The specific peptide (HDSQNLVQPGFDGMDHLITGLPFLYN) of Lhcf9 was identified. (D) Comparison of thylakoid membranes isolated under 25 °C, Air, white-light conditions and 30 °C, 3% CO_2_, white-light conditions by CN-PAGE. Thylakoid membranes were solubilized with 2% β-DDM, and the cathode buffer was supplemented with 2% Triton X-100 and 2% DOC. (E) Results of two-dimensional SDS-PAGE performed following CN-PAGE of thylakoid membranes obtained under 30 °C, 3% CO_2_, white-light conditions. (F) A representative MS/MS spectrum of the peptides of Lhcf9 in the 30 °C/3 % CO2/WL cells. The specific peptide (HDSQNLVQPGFDGMDHLITGLPFLYN) of Lhcf9 was identified.

### CgLhcf9 Expression Is Repressed by Blue Light and Enhanced under High CO_2_ Conditions

To elucidate the environmental factors regulating the expression of CgLhcf9, we examined its transcript and protein levels under different light qualities and CO_2_ concentrations using qRT-PCR and immunoblotting. We first investigated the influence of light quality. Although CgLhcf9 was previously found to accumulate under red light, it remained unclear whether this was due to a direct stimulatory effect of red light or a lack of repression by blue light. To address this, *C. gracilis* was cultured under several light conditions with varying ratios of red and blue light. Strikingly, the addition of even 1 µmol photons m^−2^ s^−1^ of blue light to red light significantly reduced CgLhcf9 expression at both RNA and protein levels compared to those under pure red-light conditions (Fig. 2A and 2D). These results indicate that CgLhcf9 expression is not directly induced by red light but is sensitively and strongly suppressed by blue light.

**Figure 2.**
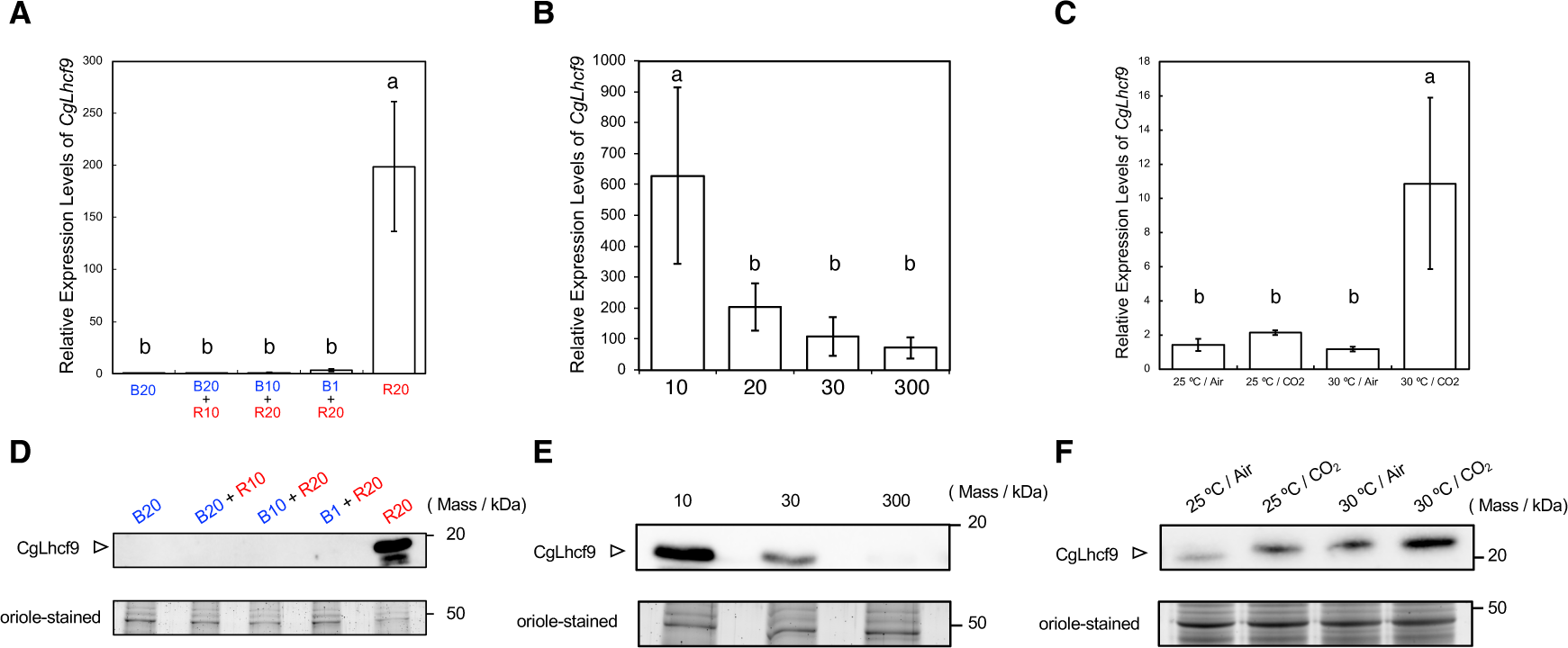
Detailed analysis of CgLhcf9 expression levels. (A–C) Relative transcript levels of CgLhcf9 were quantified by qRT-PCR from cells cultured under different conditions and are shown as means ± standard deviation (n = 3). Statistical significance was evaluated using Tukey’s HSD test (*p* < 0.05). (D–F) Immunoblot analysis of CgLhcf9 protein accumulation in whole cell extracts using anti-CgLhcf9 antibody. Equal amounts of total protein (0.3 μg per lane) were loaded for each sample. Equal loading was confirmed by Oriole-stained SDS-PAGE. (A, D) Cultures grown under different light qualities: blue light (B) and red light (R) at the indicated intensities (PPFD, µmol photons m^−2^ s^−1^). (B, E) Cultures grown under red light with varying intensities. (C, F) Cultures grown under combinations of temperature and CO_2_: 25 °C / air, 25 °C / 3% CO_2_, 30 °C / air, and 30 °C / 3% CO_2_.

Next, we assessed the red-light intensity dependence of CgLhcf9 expression. The data showed that CgLhcf9 expression was higher under low red-light intensity (Fig. 2B and 2E), suggesting that CgLhcf9 would not be involved in photoprotective mechanisms such as thermal energy dissipation. Conversely, its higher expression under low-light conditions suggests a potential role in enhancing light capture under light-limited conditions. We also evaluated its expression under four combinations of temperature and CO₂ concentration (25 °C / air, 25 °C / 3% CO_2_, 30 °C / air, and 30 °C / 3% CO_2_) with white light illumination. The qRT-PCR and immunoblotting showed that CgLhcf9 expression was induced at 25 °C with 3% CO_2_ and further enhanced at 30 °C with 3% CO_2_ (Fig. 2C and 2F).

### Generation and Spectral Characterization of CgLhcf9 Knockout and Overexpression Strains

To investigate the physiological role of CgLhcf9, we generated knockout strains (*Lhcf9-KO*) using CRISPR-Cas9 techniques and constitutive overexpression strains (*Lhcf9-OX*) that continuously express CgLhcf9. In *Lhcf9-KO* cells, no detectable expression of CgLhcf9 was observed under red light or high CO_2_ conditions (Fig. 3A and 3B). In contrast, CgLhcf9 accumulated in *Lhcf9-OX* cells even under white light with ambient air, where CgLhcf9 is not expressed in the wild-type (WT) (Fig. 3B and 3C). To examine whether CgLhcf9 expression influences the cellular absorption spectrum, we compared the spectral profiles of cells grown under various light conditions. Several eukaryotic algae, such as *Ostreobium* sp. and *Chromera velia*, are known to possess pigments and LHC proteins that are induced by far-red or red light, allowing them to effectively utilize long-wavelength light in shaded habitats, such as within coral skeletons. In such species, red light induces a shift in the absorption spectrum toward longer wavelengths, and this change is known as complementary chromatic adaptation (25–27). Since CgLhcf9 is also induced by red light, we assumed that it might play a similar role. In WT, we first confirmed that red-light-acclimated cells exhibit extended absorption beyond ∼690 nm compared to blue-light-acclimated cells, consistent with previous findings (28). However, when we compared red-light-grown *Lhcf9-KO* cells with WT, no significant difference was observed in the absorption spectrum (Fig. 3D and 3E). Similarly, under white light, the absorption spectrum of *Lhcf9-OX* cells did not exhibit a notable extension into the longer wavelengths compared to WT (Fig. 3F). These results suggest that CgLhcf9 alone does not contribute directly to the extended absorption in the long-wavelength region.

**Figure 3.**
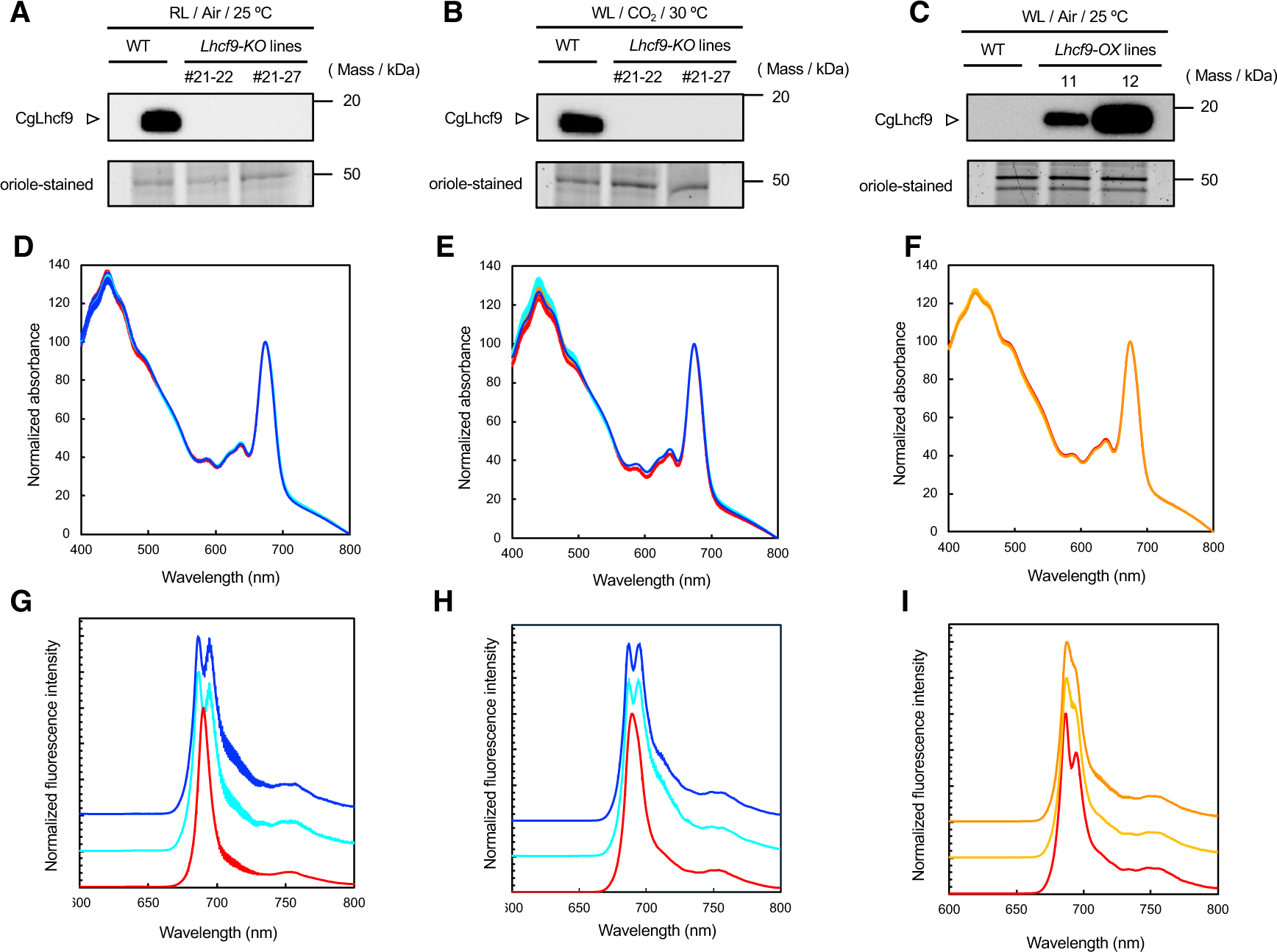
Generation and spectroscopic characterization of CgLhcf9 knockout and overexpression strains. (A–C) Immunoblot analysis of crude protein extracts from wild-type (WT), CgLhcf9 knockout (*Lhcf9-KO*), and overexpression (*Lhcf9-OX*) strains using anti-CgLhcf9 antibody. Equal amounts of total protein (0.3 µg) were loaded per lane, and Oriole-stained SDS-PAGE gels were used as a loading control. (A) WT and *Lhcf9-KO* strains cultured under red light (30 µmol photons m^−2^ s^−1^). (B) WT and *Lhcf9-KO* strains cultured under white light (30 µmol photons m^−2^ s^−1^) with 3% CO_2_. (C) WT and *Lhcf9-OX* strains cultured under white light with ambient air. (D–F) Absorption spectra of intact cells at room temperature, normalized to the Qy peak of chlorophyll *a* (∼675 nm). (D) WT (red line) and *Lhcf9-KO* (light blue (#21-22) and blue (#21-27) lines) strains under red light. (E) WT (red line) and *Lhcf9-KO* (light blue and blue lines) strains under white light with 3% CO_2_. (F) WT (red line) and *Lhcf9-OX* (yellow (OX11) and orange (OX12) lines) strains under white light with ambient air. (G–I) Steady-state fluorescence emission spectra at 77 K of intact cells excited at 459 nm. Spectra were normalized to their respective maxima. (G) WT (red line) and *Lhcf9-KO* (light blue and blue lines) strains under red light. (H) WT (red line) and *Lhcf9-KO* (light blue and blue lines) strains under white light with 3% CO_2_. (I) WT and *Lhcf9-OX* (yellow and orange lines) strains under white light with ambient air. Data in (D–I) are presented as mean ± SD from three biological replicates (n = 3).

We next examined the impact of CgLhcf9 expression on 77 K fluorescence emission spectra. In WT cultured under white or blue light, two peaks corresponding to CP43 and CP47 of PSII are typically observed at ∼686 nm and ∼694 nm, respectively (29, 30). Under red light or under high CO_2_ conditions (30 °C, white light), however, only a single peak around ∼690 nm has been reported (28). In *Lhcf9-KO* strains grown under red light or high CO_2_ conditions, this single peak was not observed; instead, the original dual peaks at ∼686 nm and ∼694 nm remained, indicating that CgLhcf9 is responsible for the ∼690 nm fluorescence peak (Fig. 3G and 3H). In *Lhcf9-OX* cells cultured under white light, although a distinct single peak at ∼690 nm was not observed, the “valley” of the fluorescence emission spectra between the CP43 and CP47 peaks in WT became less pronounced due to the fluorescence increase around ∼690 nm (Fig. 3I). This change should be caused by CgLhcf9 accumulation, which aligns with the single peak observed in red-light-acclimated WT cells. These data suggest that CgLhcf9 expression significantly alters the energy transfer process around PSII at 77 K.

### CgLhcf9 Expression Suppresses Non-Photochemical Quenching (NPQ) and Affects Cell Growth

To evaluate the impact of CgLhcf9 expression on the light-harvesting properties, we analyzed chlorophyll fluorescence, particularly focusing on the qE component of NPQ. Under red light cultivation, where WT accumulates CgLhcf9, the *Lhcf9-KO* strains consistently exhibited elevated NPQ compared to the WT (Fig. 4A). Similarly, under white light with high CO_2_ conditions, where WT accumulates CgLhcf9, NPQ levels were significantly higher in the *Lhcf9-KO* strains than in WT (Fig. 4B). Conversely, in *Lhcf9-OX* strains grown under white light, NPQ was notably reduced relative to WT, which does not accumulate CgLhcf9 (Fig. 4C). These results suggest that CgLhcf9 expression negatively regulates the qE-NPQ.

**Figure 4.**
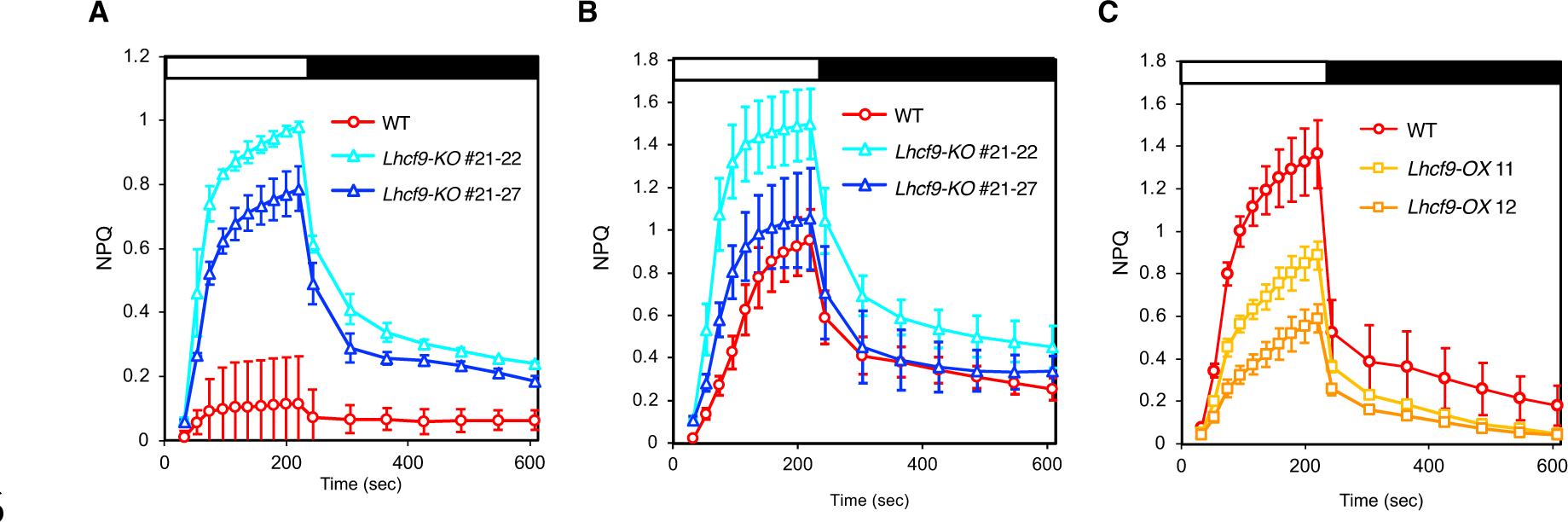
NPQ dynamics measured using chlorophyll fluorescence. Chlorophyll fluorescence was measured at room temperature using AquaPen AP-110 following 5 min dark adaptation, with excitation at 455 nm. Actinic light was set at 300 µmol photons m^−2^ s^−1^, saturation pulse at 900 µmol photons m^−2^ s^−1^, and measuring light at 0.018 µmol photons m^−2^ s^−1^. NPQ values were calculated as NPQ = (Fm − Fm′) / Fm′. Data represent means ± SD from three independent biological replicates (n = 3). (A) NPQ kinetics of wild-type (red line) and *lhcf9-KO* strains (light blue and blue lines) grown under red light (30 µmol photons m^−2^ s^−1^). (B) NPQ kinetics of wild-type (red line) and *lhcf9-KO* strains (light blue and blue lines) grown under white light (30 µmol photons m^−2^ s^−1^) with 3% CO_2_ at 30 °C. (C) NPQ kinetics of wild-type (red line) and *Lhcf9-OX* strains (yellow and orange lines) grown under white light (30 µmol photons m^−2^ s^−1^) with air.

To assess the effect of CgLhcf9 expression on cell growth, we compared the growth rates of WT and *lhcf9-KO* strains under red light at different light intensities. Under low red-light conditions (10 µmol photons m^−2^ s^−1^), WT cells, which accumulate CgLhcf9, exhibited significantly higher growth than the *lhcf9-KO* strains (Fig. 5A), while under moderate red-light conditions (30 µmol photons m^−2^ s^−1^), no marked difference in growth was observed between WT and *Lhcf9-KO* strains (Fig. 5B). In contrast, under high red-light conditions (300 µmol photons m^−2^ s^−1^), the *Lhcf9-KO* strains with higher NPQ capacities grew more rapidly than WT (Fig. 5C). We next examined growth under white light with ambient air using the *Lhcf9-OX* strains. At low light intensity (10 µmol photons m^−2^ s^−1^), *Lhcf9-OX* lines with lower NPQ showed better growth than WT (Fig. 5D). At moderate light intensity (30 µmol photons m^−2^ s^−1^), differences in growth between WT and *Lhcf9-OX* lines were minor (Fig. 5E). and no significant difference in growth was observed under high-light conditions (300 µmol photons m^−2^ s^−1^) (Fig. 5F). Furthermore, under the combined condition of 30 µmol photons m^−2^ s^−1^ white light, 30 °C, and 3% CO_2_, WT cells with higher *CgLhcf9* expression showed enhanced growth compared to the *Lhcf9-KO* strains (Fig. 5G). Collectively, these results indicate that Lhcf9 expression promotes growth under light-limiting conditions in which NPQ is less advantageous.

**Figure 5.**
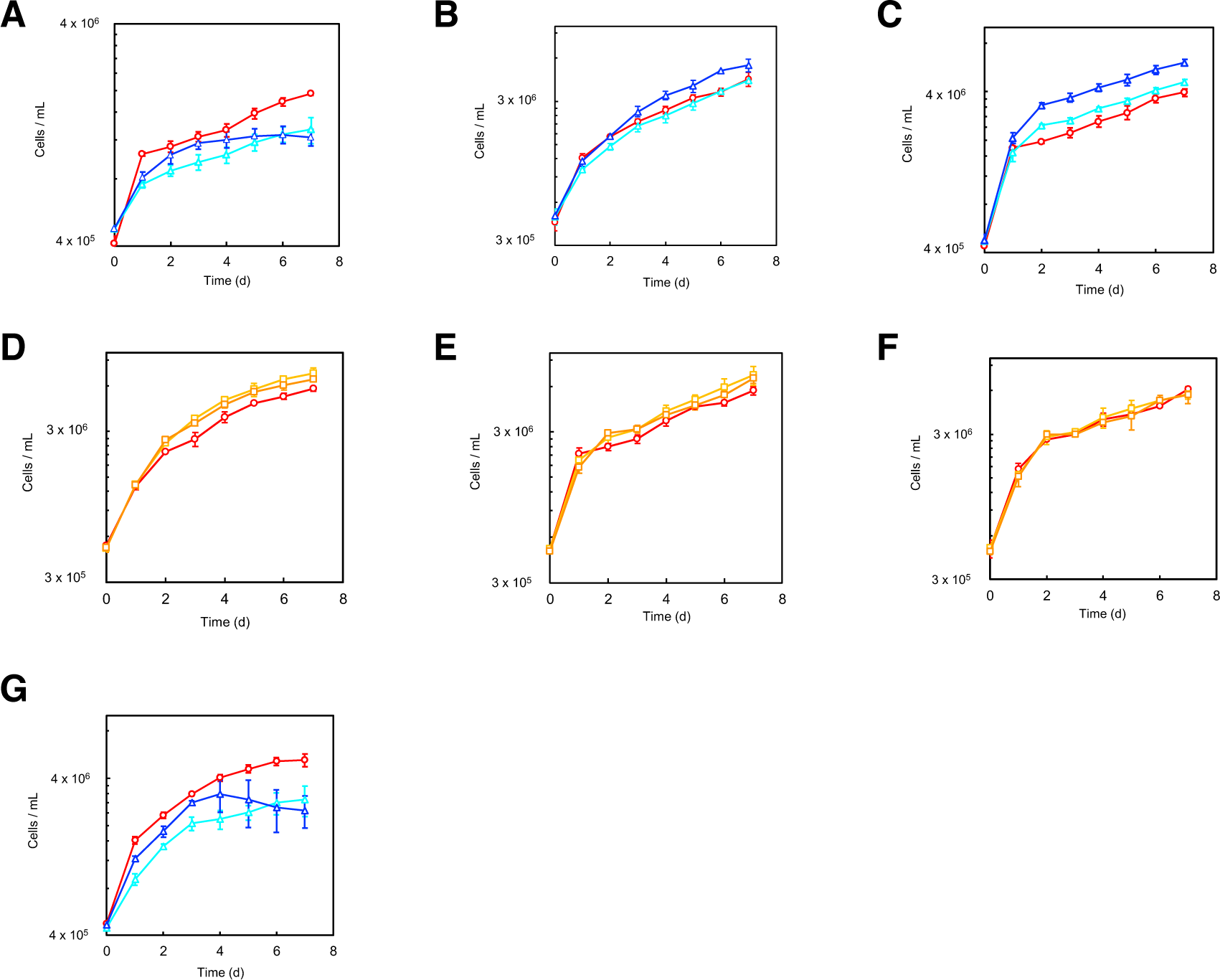
The expression of CgLhcf9 affects growth under different conditions. (A–C) Growth curves of wild-type (WT, red line) and *Lhcf9-KO* strains (light blue and blue lines) cultured under red light at low (10 µmol photons m^−2^ s^−1^; A), moderate (30 µmol photons m^−2^ s^−1^; B), and high (300 µmol photons m^−2^ s^−1^; C) intensities. (D–F) Growth curves of WT (red line) and *Lhcf9-OX* strains (yellow and orange lines) under white light at low (10 µmol photons m^−2^ s^−1^; D), moderate (30 µmol photons m^−2^ s^−1^; E), and high (300 µmol photons m^−2^ s^−1^; F) intensities. (G) Growth curves of WT (red line) and *Lhcf9*-knockout strains (light blue and blue lines) cultured under moderate white light (30 µmol photons m^−2^ s^−1^) at 30 °C with 3% (v/v) CO_2_. Cell density (cells / mL) was monitored daily, and data are presented as means ± standard deviation (n = 3, biological replicates).

### Reduction of NPQ by CgLhcf9 Is Not Attributable to Changes in Pigment Conversion nor CgLhcx1 Accumulation

Given the observed association between CgLhcf9 expression and the reduction of NPQ, we investigated whether CgLhcf9 affects the Ddx-Dtx cycle and the expression of Lhcx, both of which are known to contribute to NPQ regulation. Under low-light or dark conditions, Ddx accumulates, while under high-light conditions, Ddx is de-epoxidized to Dtx, which enhances thermal dissipation of excess excitation energy. To assess this, we calculated the Dtx/(Ddx + Dtx) ratio based on pigment quantification by HPLC. No significant difference in this ratio was observed between WT and the *Lhcf9-KO* strain under red-light conditions (Fig. 6A, Supplemental Table S2). Under white light with elevated CO_2_, a slight increase in the ratio was seen in *Lhcf9-KO* strains (Fig. 6B, Supplemental Table S3), which can be relevant to the enhanced NPQ observed. However, under white-light/Air conditions, no significant difference was observed between WT and *Lhcf9-OX* strains (Fig. 6C, Supplemental Table S4), whereas *Lhcf9-OX* strains exhibited a lower NPQ level. These results suggest that the CgLhcf9-dependent suppression of NPQ would not primarily involve alternation in the diadinoxanthin cycle.

**Figure 6.**
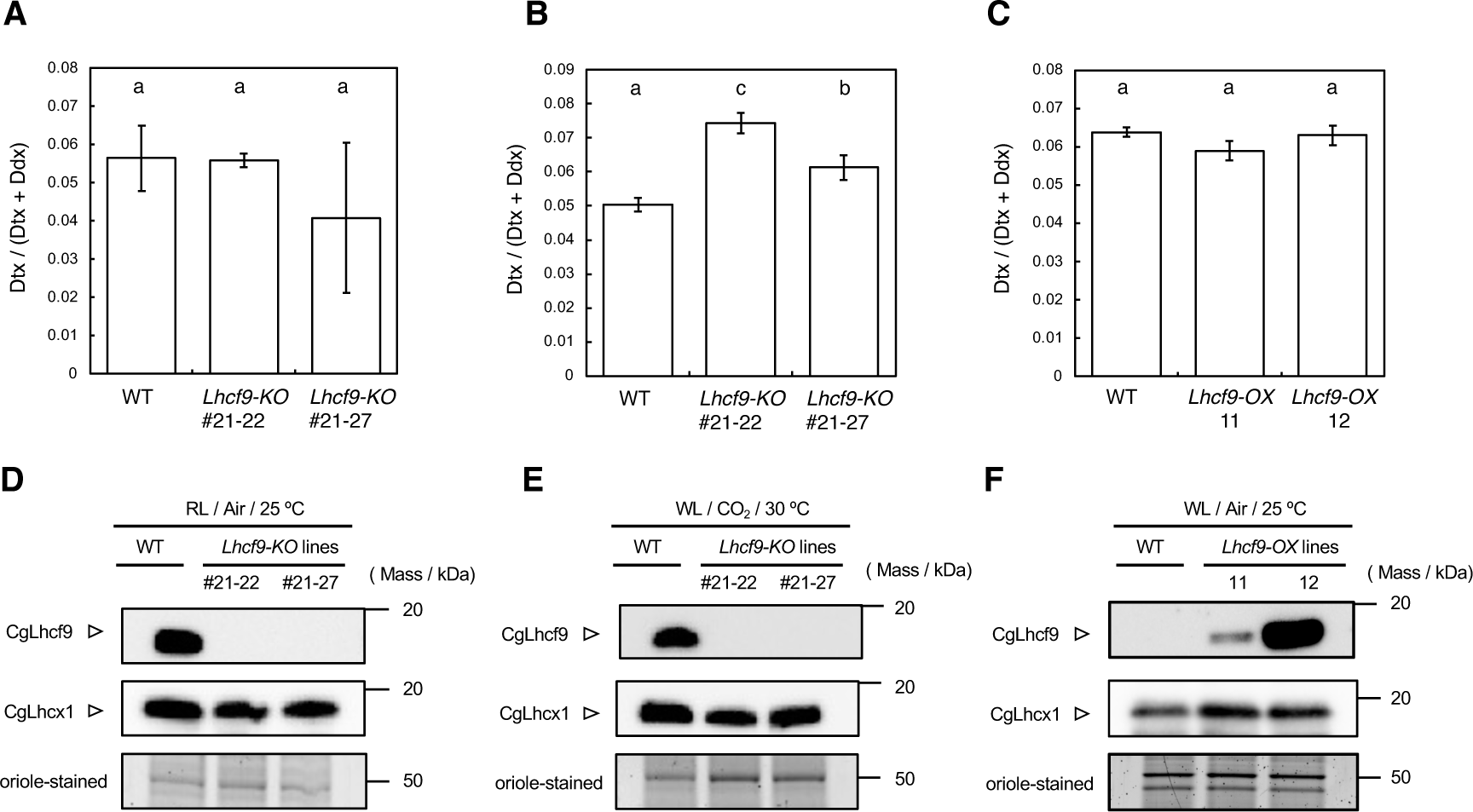
Effects of CgLhcf9 knockout and overexpression on the diadinoxanthin cycle and CgLhcx1 accumulation. (A–C) De-epoxidation state of the xanthophyll cycle, expressed as Dtx/(Ddx + Dtx), in wild-type (WT), *Lhcf9-KO* and *Lhcf9-OX* strains cultured under various conditions. (A) Red light (30 µmol photons m^−2^ s^−1^), (B) white light with 3% CO₂ at 30 °C, and (C) white light (30 µmol photons m^−2^ s^−1^) with ambient air. Values represent the mean ± SD (n = 3 biological replicates). Statistical significance was determined using Tukey’s HSD test (*p* < 0.05) and indicated by different letters. Quantitative pigment data are available in Supplementary Table 2. (D–F) Oriole-stained SDS-PAGE and immunoblot analyses of CgLhcf9 and CgLhcx1 protein levels in the corresponding culture conditions. Equal amounts of total protein (0.3 µg) were loaded for each sample. (D) WT and *Lhcf9-KO* strains under red light, (E) WT and *Lhcf9-KO* under white light with 3% CO_2_ at 30 °C, and (F) WT and *Lhcf9-OX* strains under white light with ambient CO_2_.

In *C. gracilis*, CgLhcx1 protein plays a major role in NPQ activation under excessive light stress (14). To assess whether the reduction in NPQ by CgLhcf9 is linked to changes in CgLhcx1 expression, we performed immunoblot analyses to determine protein accumulation levels of both CgLhcf9 and CgLhcx1. Under red-light conditions, CgLhcf9 was detected in WT but absent in the *Lhcf9-KO* strains. Despite the lower NPQ in WT, CgLhcx1 accumulated at comparable or even higher levels in WT than in *Lhcf9-KO* strains (Fig. 6D). A similar trend was observed under high-CO_2_ cultivation: CgLhcf9 was present in WT but absent in the *Lhcf9-KO* strains, yet CgLhcx1 was still abundant in WT, where NPQ was reduced (Fig. 6E). Under white-light/air conditions, CgLhcf9 was undetectable in WT but highly expressed in *Lhcf9-OX* strains; under this condition, CgLhcx1 accumulation remained high in *Lhcf9-OX* strains, which exhibited low NPQ, and may even have increased relative to WT. These data suggest that CgLhcf9 downregulates NPQ even in the presence of Lhcx1.

To investigate the localization of CgLhcf9 and CgLhcx1 in thylakoid membranes, CN-PAGE followed by immunoblotting with anti-CgLhcf9 and anti-CgLhcx1 antibodies was performed using the *Lhcf9-OX* strains grown under white light, which show high accumulation of both CgLhcx1 and CgLhcf9. In WT cells grown under white light, CgLhcx1 was distributed across a range of protein complex sizes, from small to large, from right to left in CN-PAGE (Fig. 7A), consistent with the report suggesting its potential interaction with oligomeric LHC complexes (14). In contrast, in the *Lhcf9-OX* strain, in which CgLhcf9 accumulated at higher levels, the broad distribution of CgLhcx1 appeared to be shifted toward the right, smaller protein region (Fig. 7B). These trends were also observed when immunoblotting was performed directly after CN-PAGE (Supplemental Fig. S7). Interestingly, CgLhcf9 exhibited a similar distribution pattern to CgLhcx1 in WT thylakoids, spanning smaller to larger protein regions in CN-PAGE. This suggests the possibility that CgLhcf9 accumulation may affect the association of CgLhcx1 to the peripheral LHC complexes, probably loosely associated with PSII–FCPII, preventing the formation of energy quenching sites for the qE-NPQ.

**Figure 7.**
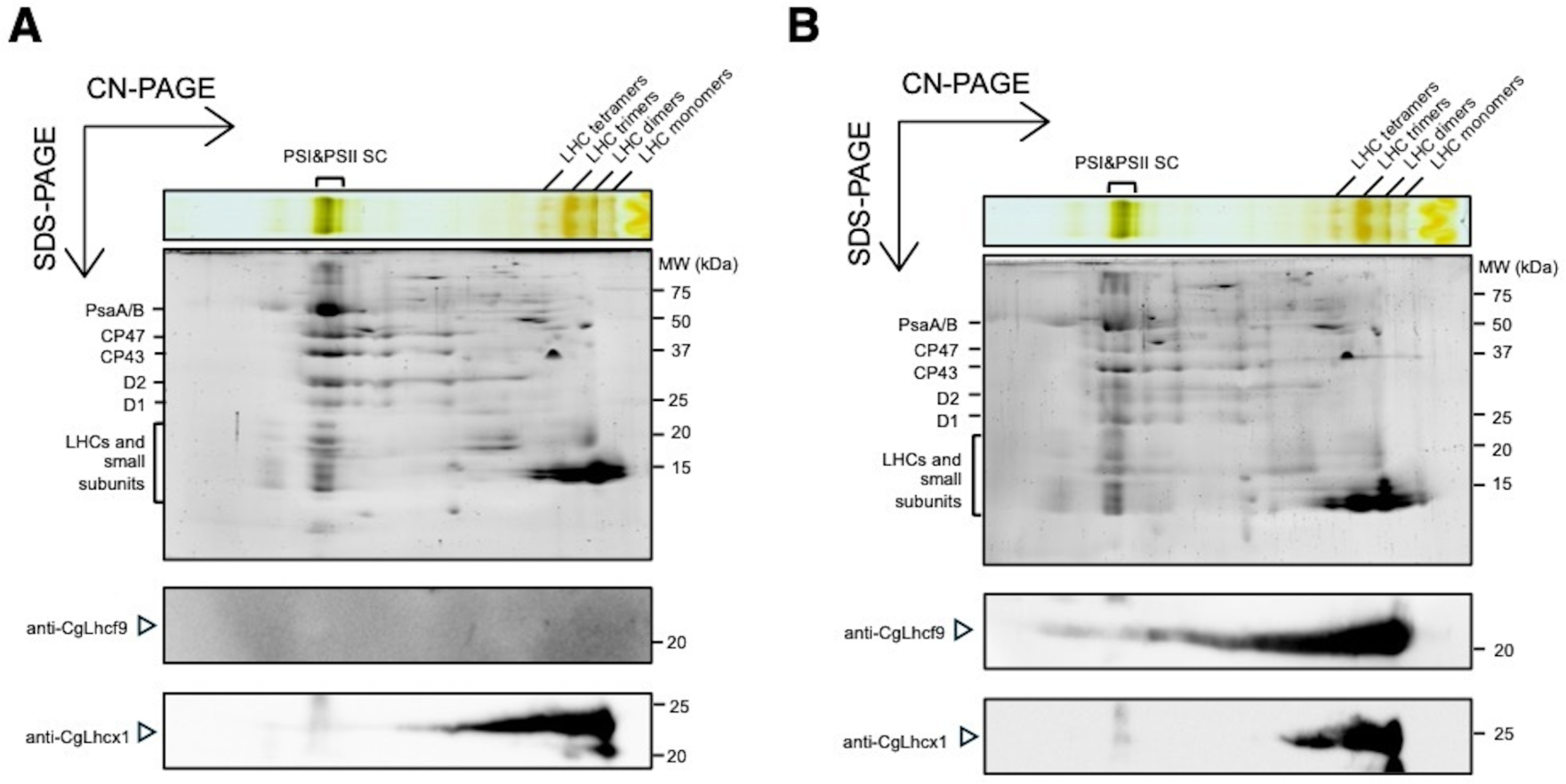
Localization of CgLhcf9 and CgLhcx1. (A), (B) CN-PAGE and 2D SDS-PAGE followed by immunoblotting with anti-CgLhcx1 and anti-CgLhcf9 antibodies. (A) WT under white light condition with ambient air. (B) *Lhcf9-OX* line under white light condition with ambient air.

## Discussion

Aquatic light environments are highly dynamic, requiring photosynthetic algae to develop flexible adaptive strategies for survival. Among these strategies, various light-harvesting complex (LHC) proteins, also known as fucoxanthin chlorophyll *a*/*c*-binding proteins (FCPs) in diatoms, play a crucial role in adjusting to fluctuating light conditions. In this study, we focused on CgLhcf9, a unique LHC that is specifically induced under red light and high CO_2_ conditions (Fig. 1). CgLhcf9 displays a distinct gene expression profile, markedly different from other LHC proteins, suggesting a potential role in optimizing photosynthesis in response to environmental stimuli.

Detailed expression analyses revealed that CgLhcf9 is strongly suppressed by blue light (Fig. 2A and 2B), and its expression increases under low-intensity red light (Fig. 2C and 2D). Blue light is recognized as a major inducer of NPQ in both green algae and diatoms (31–34). In the model diatom *P. tricornutum*, Aureochrome1c (Aureo1c)̶a heterokont-specific photoreceptor with a blue-light-sensing LOV domain̶has been identified as a key regulator of NPQ. Aureo1c also has a bZIP DNA-binding domain and functions as a transcription factor, directly activating NPQ-related *Lhcx* genes: It binds the promoter regions of the NPQ-inducing factors *Lhcx2* and *Lhcx3*, and its disruption results in decreased NPQ (35). In contrast to the above blue-light-induced NPQ factors, CgLhcf9 is suppressed by blue light, suggesting that CgLhcf9 functions in light harvesting rather than in NPQ. Additionally, we observed that transcription of the *CgLhcf9* gene was enhanced under 30 °C / 3% CO_2_ condition (Fig. 2C), highlighting its multifaceted environmental responsiveness.

NPQ is typically associated with the dissipation of excess light energy, while recent studies in green algae have revealed its broader physiological relevance. In particular, Lhcsr—the key protein responsible for NPQ in *Chlamydomonas reinhardtii*—is downstream of the CO_2_-concentrating mechanism (CCM) signaling pathway, responding not only to light intensity but also to changes in CO_2_ availability to regulate NPQ (36). This NPQ regulation via Lhcsr helps balance the light-dependent electron transport reactions with CO_2_ fixation. For example, under low CO_2_ conditions, even moderate light levels can lead to an over-reduction of the photosynthetic electron transport chain, due to the limited electron sinks resulting from reduced carbon fixation. This situation necessitates the activation of NPQ by Lhcsr as a photoprotective response. In this way, Lhsr triggers NPQ not only under strong or blue light but also in response to metabolic imbalances caused by limited CO_2_ supply. These dual regulatory pathways via Lhcsr in green algae highlight NPQ’s crucial role in enhancing photosynthetic efficiency under various environmental conditions.

In diatoms, this coordinated regulation between excitation pressure and electron sink may rely on distinct mechanisms, because their CCM has been acquired convergently and independently from green algae (37, 38). We propose that CgLhcf9 is the key component for this fine-tuning: CgLhcf9 expression was markedly induced under red light, particularly at low light intensities, whereas it was strongly suppressed in the presence of blue light. In addition, its accumulation was also enhanced under elevated CO_2_ conditions where there is plenty of electron sinks (Fig. 2). When these expression patterns are compared with the conditions known to promote NPQ—namely, high light (especially blue light) and low CO_2_ availability—it becomes evident that CgLhcf9 is upregulated under environmental conditions where NPQ is not induced. This inverse relationship indicates that CgLhcf9 can be a negative regulator of NPQ, suppressing NPQ when photoprotection is not required.

This hypothesis is further supported by functional analysis using *Lhcf9* knockout (*lhcf9-KO*) and overexpression (*Lhcf9-OX*) strains. Under red light conditions, the *Lhcf9-KO* strains exhibited a significant increase in NPQ compared to the wild type (WT) (Fig. 4A). Similarly, under high CO_2_ conditions, *Lhcf9-KO* strains showed elevated NPQ relative to WT (Fig. 4C). In contrast, under white light with ambient air, *Lhcf9-OX* strains displayed reduced NPQ compared to WT (Fig. 4E). These results strongly support that CgLhcf9 acts as a negative regulator of NPQ, functioning to limit energy dissipation under conditions where photoprotection is less critical.

The appearance of a single fluorescence peak at ∼690 nm between the CP43 (∼686 nm) and CP47 (∼696 nm) peaks in the 77 K fluorescence emission spectra of strains expressing CgLhcf9 suggests the alternation of the energy transfer to the PSII core antenna (Fig. 3G-I). The presence of a single distinct peak implies that CgLhcf9 would serve as an energy trap for excitation energy at cryogenic temperatures. Given that CgLhcf9 is not identified in the PSII-LHCII supercomplex in CN-PAGE, it suggests that a tight complex with PSII and LHCII is not changed even under CgLhcf9-expressing conditions, and CgLhcf9 serves as the peripheral antenna of PSII-LHCII. The different excitation energy transfer at 77 K suggests the different kinetics of excitation energy transfer even at ambient temperatures. This alteration in excitation energy flow would contribute to the observed suppression of NPQ, suggesting that CgLhcf9 may play a structural and functional role in modulating energy transfer pathways to the PSII core.

Analysis using both *Lhcf9-KO* and *Lhcf9-OX* strains revealed that CgLhcf9 alters NPQ without affecting the amount of CgLhcx and Dtx (Fig. 6A-C). CgLhcx1 seems to interact with peripheral LHC complexes as indicated by immunoblotting following CN/SDS-PAGE, showing that CgLhcx1 exhibits a broad distribution extending from smaller to larger protein-complex regions (14). The induction of energy quenching in qE at the peripheral antenna by CgLhcx1 is also supported by time-resolved fluorescence analysis (14). Furthermore, CgLhcx1 becomes unstable in the mutant lacking CgLhcf2, one of the peripheral LHC complexes surrounding PSII-LHCII supercomplex (39). In *Lhcf9-OX* cells, the broad distribution of CgLhcx1 appeared to be shifted toward smaller protein complex, and the band pattern of CgLhcf9 in *Lhcf9-OX* strains resembled that of CgLhcx1 observed in WT cells under white light (Fig. 7A-B). These facts suggest that CgLhcf9 would alter the association of CgLhcx with the peripheral LHC complexes (Figure 8). CgLhcf9 may interact with CgLhcx1; However, the interaction must be weak or transient because we have not been successful in isolating the CgLhcx1-CgLhcf9 complex.

**Figure 8.**
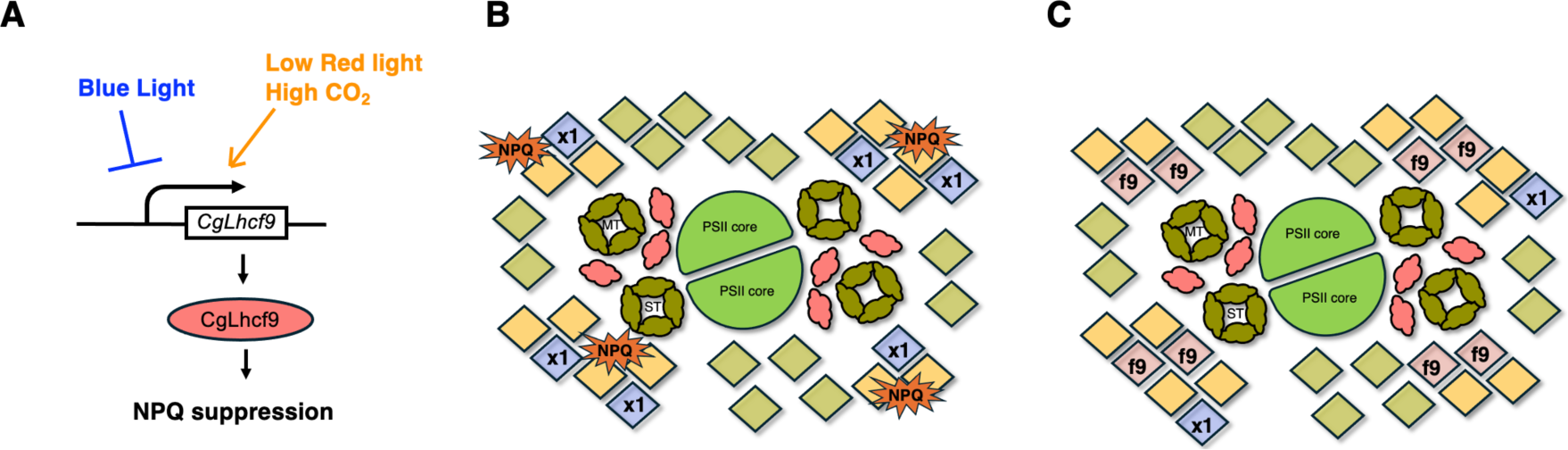
A hypothetical model for negative regulation of qE-NPQ via CgLhcf9. (A) CgLhcf9 expression is down-regulated by blue light and up-regulated by low-red light and high CO₂ conditions, which are opposite to the environmental factors inducing qE-NPQ. The accumulation of CgLhcf9 suppresses qE-NPQ, indicating that CgLhcf9 is a novel negative regulator of energy dissipation in the marine diatom *C. gracilis*. (B) Lhcx1 (x1) associates with peripheral LHCs around PSII-LHCII complex to induce NPQ. (C) Accumulation of Lhcf9 (f9) affects the interaction between Lhcx1 and peripheral LHCs, resulting in the exclusion of Lhcx1and a decreased NPQ.

Overall, our findings suggest that CgLhcf9 expression is regulated in response to environmental factors, such as light quality and CO_2_ concentration, thereby modulating NPQ and contributing to the optimization of the photosynthetic energy conversion. Such plastic changes in the antenna system would be a key adaptive mechanism to maintain fitness under fluctuating environmental conditions. Phylogenetic analysis indicates homologs for CgLhcf9 in *P. tricornutum* (21). Among them, PtLhcf15 shows a ∼10-fold increase in expression under red light and emits a characteristic long-wavelength fluorescence at both room temperature and 77 K, known as F710 (3, 24, 40). Unlike PtLhcf15, CgLhcf9 does not emit a far-red fluorescence and displays a single fluorescence peak at ∼690 nm between CP43 (∼686 nm) and CP47 (∼696 nm) peaks at 77K. Although F710 emission is often regarded as a hallmark of long-wavelength adaptation, such a feature is not observed in the case of CgLhcf9, indicating a different functional property. Noticeably, CgLhcf9 is phylogenetically closer to PtLhcf13 and has not been identified as a component of the F710-emitting complex, although it is upregulated by approximately 20% under red light (3, 24). Further experimental validation is needed to clarify whether the red light-induced LHCs negatively regulate NPQ in other diatom species.

## Materials and Methods

### Cell Culture

All experiments were performed using the marine centric diatom *Chaetoceros gracilis* (UTEX LB 2658). Cells were grown in artificial seawater prepared with 4% (w/v) Daigo artificial sea salts supplemented with 1× Daigo IMK medium (Nihon Pharmaceutical, Japan) and 0.2 mM Na_2_SiO_3_. For preculturing, cells were inoculated under continuous shaking (100 rpm) at 25 °C with white light (30 µmol photons m^−2^ s^−1^). For main cultures, cells were inoculated at an initial density of 5 × 10^5^ cells/mL and grown under white, red (660 nm), or blue (450 nm) LED light with continuous aeration using either ambient air or 3% (v/v) CO_2_.

Genetic transformation was conducted via electroporation using the NEPA21 Type II Super Electroporator (NEPAGENE, Japan) following previously established protocols for *C. gracili*s transformation with hCas9 and target genes (14, 41–43). Cell growth was monitored using a Luna™ Automated Cell Counter (Logos Biosystems). Chlorophyll content was determined in 90% acetone based on Jeffrey & Humphrey (1975).

### RNA Extraction and Quantitative RT-PCR

Total RNA was extracted from *C. gracilis* cells using the RNeasy Plant Mini Kit (Qiagen Inc., Valencia, CA, USA), with modifications to the manufacturer’s protocol based on a previously reported method (21). Genomic DNA was removed, and cDNA synthesis was performed using the PrimeScript™ RT reagent Kit with gDNA Eraser (Perfect Real Time; Takara Bio, Japan) according to the manufacturer’s instructions. Quantitative real-time PCR was carried out using a CFX96™ Real-Time System (Bio-Rad Laboratories, Hercules, CA, USA).

### SDS-PAGE and Immunoblotting

Total protein was extracted from harvested cells by solubilization in sample buffer (62.5 m mol L^−1^ Tris–HCl pH 6.8, 2.5% SDS, 10% glycerol, 2.5% 2-mercaptoethanol, trace bromophenol blue), followed by incubation at 37 °C for 30 min and centrifugation. Protein concentrations were determined using the RC DC Protein Assay Kit (Bio-Rad), and samples were adjusted to 3 μg/10 μL. Proteins were separated by SDS-PAGE using a 16% polyacrylamide gel containing 6 mol L^−1^ urea under 250 V, 50 mA for 90 min. After electrophoresis, gels were stained with Oriole™ fluorescent stain for 90 min and imaged using the ChemiDoc Touch Imaging System (Bio-Rad, CA, USA).

For Western blotting, proteins were transferred to a PVDF membrane (Immobilon-P, Millipore, 0.45 μm) using the Trans-Blot Turbo system (Bio-Rad, CA, USA). Anti-CgLhcf9 antibody (1:4000) was used. Blocking, primary and secondary antibody incubations were performed using Can Get Signal solution (Toyobo, Japan). Detection was conducted with Amersham™ ECL Prime reagent (Cytiva), and signals were visualized using the ChemiDoc Touch Imaging System (Bio-Rad, CA, USA).

### 2D-CN/SDS-PAGE and Mass Spectrometry

Thylakoid membranes were isolated as previously described (14). Clear native (CN)-PAGE was performed using 4–13% gradient polyacrylamide gels prepared according to established protocols (14, 44, 45). Thylakoid suspensions were adjusted to 1 μg Chl μL^−1^ in solubilization buffer (50 mM imidazole-HCl, pH 7.0 at 4 °C, 20% glycerol). For amphipol-based solubilization, equal volumes of 2% (w/v) α-dodecyl maltoside (α-DDM) were added, incubated on ice for 2 min, and centrifuged at 21,500 × g for 2 min. For DOC-Triton-based CN-PAGE, solubilization was done with 2% α-DDM; for DOC-based CN-PAGE, 2% β-dodecyl maltoside or 2% TritonX-100 was used. After 30 min on ice and centrifugation (21,500 × g, 2 min), 5 μg of Chl in 10 μL was loaded. The anode buffer contained 25 mmol L^−1^ imidazole (pH 7.0), and the cathode buffer contained 50 mmol L^−1^ Tricine and 7.5 mmol L^−1^ imidazole. For DOC-Triton-based runs, the cathode buffer additionally contained 0.02% sodium deoxycholate and 0.02% Triton X-100; for DOC-based runs, 0.05% sodium deoxycholate was used.

After electrophoresis, CN-PAGE gels were soaked in 50% (v/v) glycerol on ice, cut into gel strips, and stored at −80 °C. Gel strips were incubated for 30 min at room temperature in solubilization buffer (1% SDS, 50 mmol L^−1^ DTT) and applied to second-dimension SDS-PAGE (stacking gel: pH 6.8; separation gel: 14% acrylamide, 6 mol L^−1^ urea, pH 8.6). Electrophoresis was performed using the Laemmli system. For protein visualization (when not used for immunoblotting), gels were stained for 90 min with Oriole fluorescent stain (#1610496, Bio-Rad) and imaged using the ChemiDoc Touch Imaging System (Bio-Rad).

For mass spectrometry, protein bands were excised, destained, and digested with TPCK-treated trypsin (Worthington Biochemical). Peptides were analyzed using Easy nLC 1000 coupled with a Q Exactive mass spectrometer (Thermo Fisher Scientific), and spectra were processed as described previously (18).

### Pulse-Amplitude Modulation Chl Fluorescence Analysis

Chlorophyll fluorescence was measured using a pulse-amplitude modulation (PAM) protocol (NPQ2) with an AquaPen AP-110 (PSI, Czech Republic). Cells were illuminated with actinic light at 455 nm (300 μmol photons m^−2^ s^−1^), and fluorescence emission was detected through a bandpass filter (667–750 nm). The NPQ2 protocol included a 200 s light phase followed by a 390 s dark recovery phase, with the first saturation pulse applied 10 s after the onset of actinic illumination. Measuring light intensity was set to 10%, and saturation pulse intensity was set to 30%, equivalent to 900 μmol photons m^−2^ s^−1^. Prior to measurement, cells were dark-adapted for 5 min.

### Spectroscopic Measurements

Absorption spectra were recorded using a UV-2600 spectrophotometer (Shimadzu, Japan) equipped with a photomultiplier unit. Cell suspensions (3 mL) were placed in a quartz cuvette, and absorbance was measured from 400 to 800 nm at 0.5 nm intervals.

Steady-state fluorescence spectra at 77 K were obtained using a JASCO FP-8500 spectrofluorometer equipped with a PMU-130 liquid nitrogen cooling unit. Cell suspensions were adjusted to 2 μg Chl *a*+*c*/mL and excited at 459 nm. Spectra were averaged from five successive scans and collected with a sampling pitch of 2.5 nm. Fluorescence intensities were normalized to the maximum peak.

To estimate the relative PSII antenna size, chlorophyll fluorescence induction curves were measured using a Fluorometer FL-3500 (PSI, Czech Republic) following 5 min of dark acclimation. The measurement was conducted in the presence of 40μM DCMU. Flash intensity and duration were set to 75% and 100 μs, respectively. The antenna size was evaluated based on the slope of the fluorescence rise from initial fluorescence to the level corresponding to two-thirds of the maximum intensity (46).

### HPLC Pigment Analysis

Cells were harvested by centrifugation at 1,500 × g for 5 min, and the supernatant was discarded. Cell pellets were immediately frozen in liquid nitrogen and stored at −80 °C until pigment extraction. Frozen pellets were extracted with 100% acetone (Wako, Japan) using ultrasonic disruption for 6 min in an ice-water bath. The extracts were centrifuged at 15,000 rpm at 4 °C for 5 min, and the supernatants were filtered through a 0.45 μm hydrophilic PTFE membrane filter (SLPT0445NL, Hawach Scientific, IL, USA) before HPLC analysis.

HPLC pigment analysis was conducted according to the modified method based on Zapata et al. (2000) (47), as described in Nagao et al. (2020) (23). Solvent A (containing 0.25 mol L^−1^ pyridine) was adjusted to pH 5.0 using acetic acid. Pigments were analyzed using a Shimadzu HPLC system equipped with an LC-20AD pump and SPD-M20A photodiode array detector. An Inertsil C8 reversed-phase column (5020-01228, GL Sciences, Japan) was used, and 20 μL of each sample was injected for analysis.

## Author Contributions

M.K. and K.I. conceived the project; M.N. performed growth experiments, qRT-PCR, immunoblotting, 2D-CN/SDS-PAGE, PAM analyses, absorbance spectrum analyses, 77 K emission fluorescence analysis, antenna size measurements; M.K., R.N., T.S., and N.D. performed mass spectrometry; M.N. and S.T. performed HPLC analysis; M.N., M.K., and N.I. created knockout and overexpression strains; H.H., H.T., A.S., and S.I. performed RNA-seq and provided funding; S.A. performed time-resolved fluorescence analyses; M.N. drafted the original manuscript. M.K. and K.I. revised and edited the final manuscript. All authors contributed to the discussion of the results and improvement of the manuscript.

## Acknowledgement

This work was supported in part by JSPS KAKENHI grant nos. JP24KJ1498 (M.N.), JP22KJ2017 (M.K.), JP24H02086 (S.A.), JP23K27040 (K.I. and S.A.), JP24H02081 (K.I.), and a grant no. L-2023-3-007 from the Institute for Fermentation, Osaka, Japan (K.I.).

