## Supplemental for "Lhcf9 is a novel negative regulator of non-photochemical quenching in the diatom *Chaetoceros gracilis*"

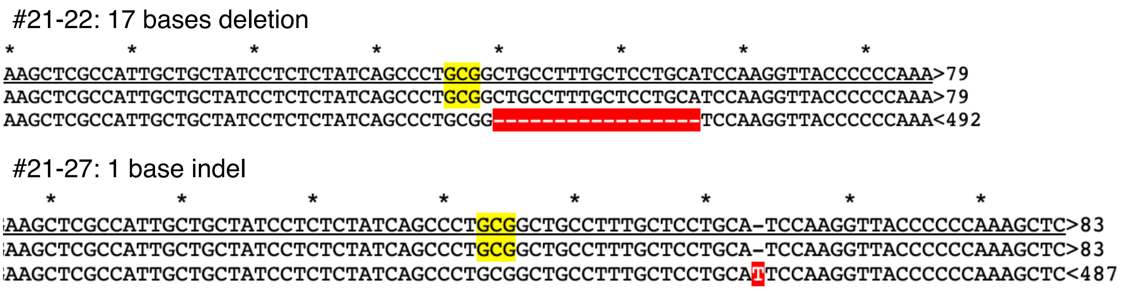


**Supplemental Figure S1. Confirmations of *Lhcf9-KO* lines**

Mutations in Lhcf9 knockout (*Lhcf9-KO*) lines generated by CRISPR–Cas9 were confirmed by Sanger sequencing.

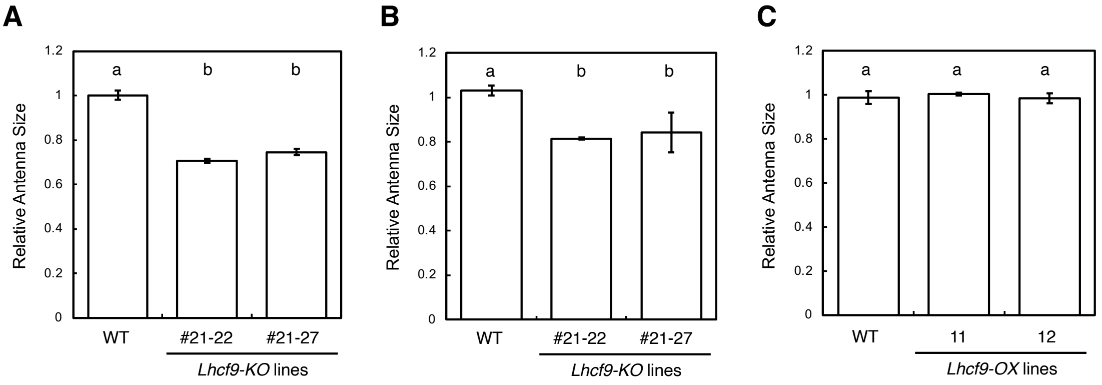


**Supplemental Figure S2. Relative PSII antenna size**

PSII antenna size was determined from the initial slope of the increase in fluorescence yield until reaching two-thirds of the maximum fluorescence intensity. Relative antenna size was normalized to the WT value, which was set to 1. Values represent means ± SD from three biological replicates (n = 3). Different letters indicate statistically significant differences as determined by one-way ANOVA followed by Tukey's HSD test ($p < 0.05$). (A) WT and *Lhcf9-KO* strains under red light. (B) WT and *Lhcf9-KO* strains under white light with 3% $CO_2$. (C) WT and *Lhcf9-OX* strains under white light with ambient air. Means ± SD from three biological replicates (n = 3).

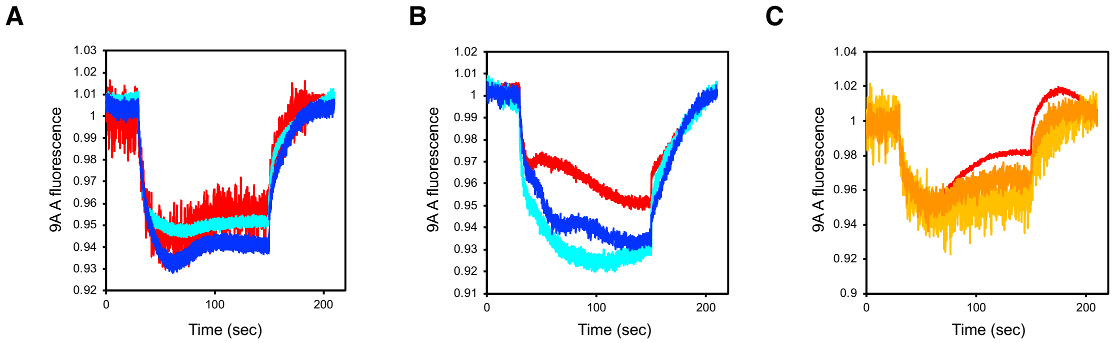


**Supplemental Figure S3. ΔpH 9-AA fluorescence**

(A)-(C) WT (red line) and *Lhcf9-KO* (light blue and blue lines) strains under red light. (B) WT (red line) and *Lhcf9-KO* (light blue and blue lines) strains under white light with 3% $CO_2$. (C) WT (red line) and *Lhcf9-OX* (yellow and orange lines) strains under white light with ambient air. Means ± SD from three biological replicates (n = 3).

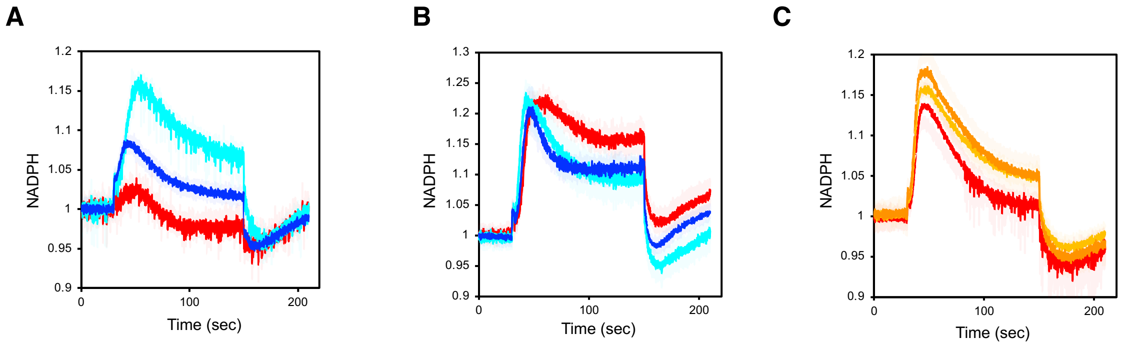


**Supplemental Figure S4. NADPH measurements**

(A)-(C) WT (red line) and *Lhcf9-KO* (light blue and blue lines) strains under red light. (B) WT (red line) and *Lhcf9-KO* (light blue and blue lines) strains under white light with 3% $CO_2$. (C) WT (red line) and *Lhcf9-OX* (yellow and orange lines) strains under white light with ambient air. Means ± standard deviation from three biological replicates (n = 3).

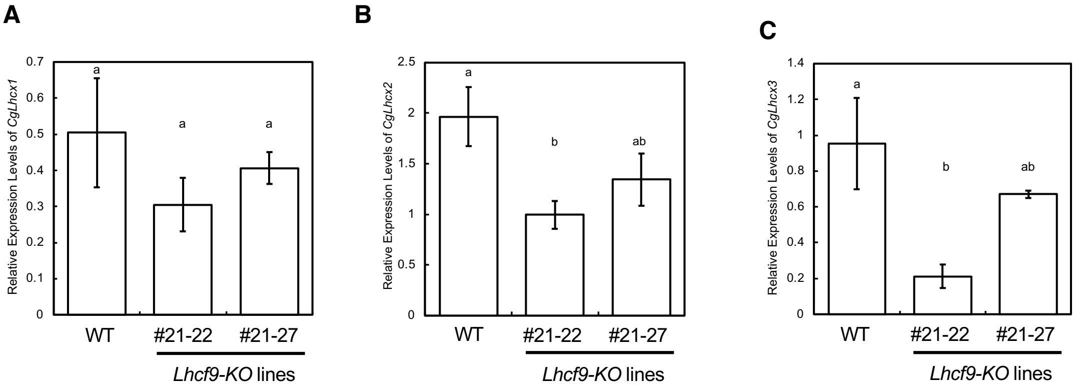


**Supplemental Figure S5. Expression levels of CgLhcx1-3**

(A)-(C) Relative transcript levels of *CgLhcx*1(A), *CgLhcx2*(B) and *CgLhcx3*(C) were quantified by qRT-PCR from cells cultured under red light conditions and are shown as means ± standard deviation (n = 3). Statistical significance was evaluated using Tukey's HSD test ($p < 0.05$).

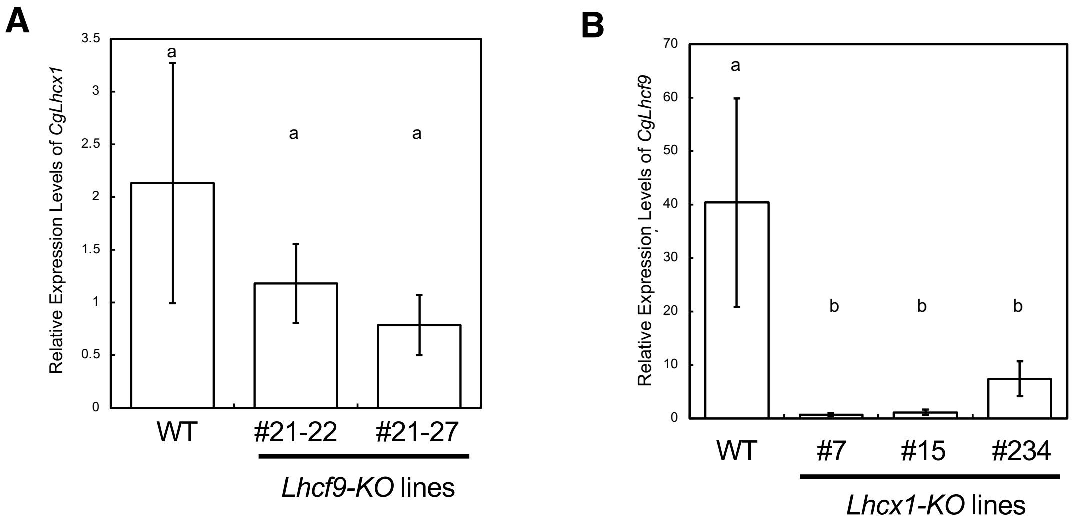


**Supplemental Figure S6. Expression relationship between *CgLhcf9* and *CgLhcx1* under high $CO_2$ growth conditions**

Relative transcript levels of *CgLhcx*1(A) and *CgLhcf9* (B) were quantified by qRT-PCR from cells cultured under high light/high $CO_2$ conditions and are shown as means ± standard deviation (n = 3). Statistical significance was evaluated using Tukey's HSD test ($p < 0.05$).

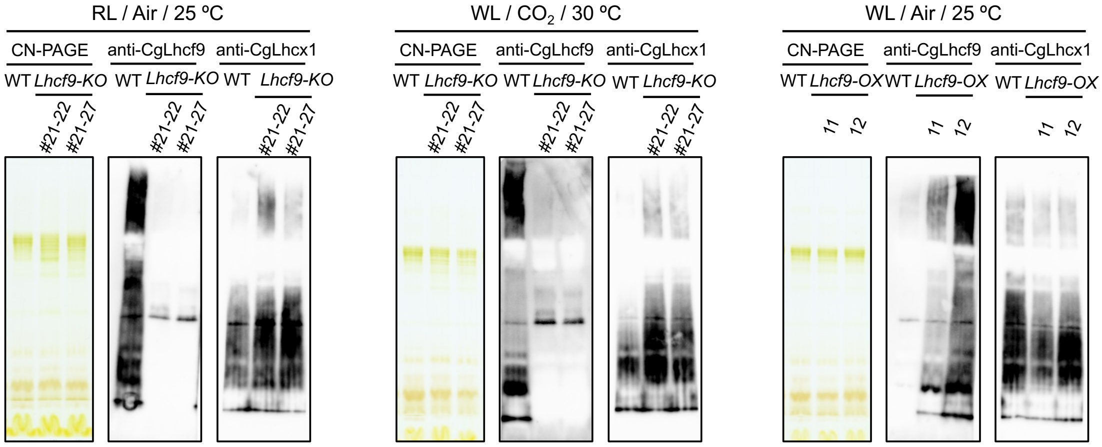


**Supplemental Figure S7. Localization of CgLhcf9 and CgLhcx1**

(A)-(C) CN-PAGE and immunoblotting with anti-CgLhcf9 and anti-CgLhcx1 antibodies of thylakoid membranes isolated from (A)WT and *Lhcf9-KO* strains under red light cultivation, (B) WT and *Lhcf9-KO* strains under white light with 3% $CO_2$ cultivation and (C) WT and *Lhcf9-OX* strains under white light with ambient air.

**Supplemental Table S1. Primers**

| Primer name | Sequence (5´ → 3´) |
|---|---|
| Lhcf9_21-43_RGR_BamHI_F1 | GTCAGGATCCaggataCTGATGAGTCCGTGAGGACGAAACGAGTAAGCTCGTC |
| Lhcf9_21-43_RGR_F2 | GACGAAACGAGTAAGCTCGTCtatcctctctatcagccctgGTTTTAGAGCTAGAAATAGCAAG |
| pCgNRp526_Fw | TTGTGCCAGAACAAAAGACAA |
| PstI_RGR_Rv | ATATCTGCAGGTCCCATTCGCCATGCCGAAGC |
| Cg Lhcf9 453bp 22011 Fw | CATGCGCATTACTGGAATTG |
| Cg Lhcf9 453bp 22011 Rv | TGTCAAAGGGGAAACGGTAG |
| 230405_BamHI_Lhcf9_for_OX_Fw | TATATAGGATCCATGAAGCTCGCCATTGCTGC |
| 230405_Lhcf9_PstI_for_OX_Rv | ATATATCTGCAGTTAATTGTAAAGGAATGGGAGCCCAGTG |
| 230405_CgLhcf4p_f4pro_Fw | ATCTTTCTGTCGTGAGCAGGA |
| 230405_CgLhcf4p_Rv | TCTGCTGAATTGGGCAATTGG |
| CgLhcf9_211222_Fw | AGATCATGAAGCTCGCCATT |
| CgLhcf9_211222_Rv | TGACACCAAGCTCATTCTCG |
| 20221026_Cg_a_tubulin_qPCR_Fw | GCTTTATCATCCCGAGCAAA |
| 20221026_Cg_a_tubulin_qPCR_Rv | GTCGCATGGAACACAAGAAA |
| 20240116_CgLhcx1_qPCR_Fw | GCCATCTCTGCTCTCCTTATT |
| 20240116_CgLhcx1__qPCR_Rv | AAGAGGATCGAAGATTCCTACA |
| 20240116_CgLhcx2_qPCR_Fw | GCTTTTATTGGAGCTTATGAGGC |
| 20240116_CgLhcx2_qPCR_Rv | CCAAAGGGGTCAAACTTGAG |
| 20240116_CgLhcx3_qPCR_Fw | AGCTCGTCCTTGAGTATGTCC |
| 20240116_CgLhcx3_qPCR_Rv | ATTGATGGGAGTGACAGCAGG |

**Supplemental Table S2. Pigments of WT and *Lhcf9-KO* lines under red light condition**

Fucoxanthin (Fx), diadinoxanthin (Ddx), diatoxanthin (Dtx), and β-carotene (β-Car) contents were quantified by HPLC and expressed as molar ratios to chlorophyll *a* (Chl *a*) (mol/mol). de-epoxidation state of the diadinoxanthin cycle is shown as Dtx/ (Ddx + Dtx), calculated on a molar basis. Values are means ± standard deviation from biological replicates (n = 3).

| Sample (RL) | Fx / Chl *a* | Ddx / Chl *a* | Dtx / Chl *a* | Dtx / (Ddx + Dtx) | β-Car / Chl *a* |
|---|---|---|---|---|---|
| WT | 0.66 ± 0.037 | 0.22 ± 0.016 | 0.014 ± 0.0029 | 0.056 ± 0.0086 | 0.040 ± 0.0026 |
| *Lhcf9-KO* #21-22 | 0.63 ± 0.022 | 0.26 ± 0.010 | 0.015 ± 0.00065 | 0.056 ± 0.0018 | 0.057 ± 0.0016 |
| *Lhcf9-KO* #21-27 | 0.66 ± 0.027 | 0.22 ± 0.0042 | 0.0096 ± 0.0049 | 0.041 ± 0.020 | 0.048 ± 0.00040 |

**Supplemental Table S3. Pigments of WT and *Lhcf9-KO* lines under WL / 3 % CO2 / 30 ºC condition**

Fucoxanthin (Fx), diadinoxanthin (Ddx), diatoxanthin (Dtx), and β-carotene (β-Car) contents were quantified by HPLC and expressed as molar ratios to chlorophyll *a* (Chl *a*) (mol/mol). de-epoxidation state of the diadinoxanthin cycle is shown as Dtx/ (Ddx + Dtx), calculated on a molar basis. Values are means ± standard deviation from biological replicates (n = 3).

| Sample ($CO_2$) | Fx / Chl *a* | Ddx / Chl *a* | Dtx / Chl *a* | Dtx / (Ddx + Dtx) | β-Car / Chl *a* |
|---|---|---|---|---|---|
| WT | 0.75 ± 0.0084 | 0.26 ± 0.0046 | 0.0014 ± 0.00081 | 0.050 ± 0.0020 | 0.046 ± 0.0016 |
| *Lhcf9-KO* #21-22 | 0.67 ± 0.0099 | 0.35 ± 0.016 | 0.028 ± 0.0026 | 0.074 ± 0.0030 | 0.059 ± 0.0039 |
| *Lhcf9-KO* #21-27 | 0.70 ± 0.016 | 0.27 ± 0.0032 | 0.017 ± 0.0013 | 0.061 ± 0.0037 | 0.052 ± 0.0025 |

**Supplemental Table S4**. **Pigments of WT and *Lhcf9-OX* lines under WL / Air condition**

Fucoxanthin (Fx), diadinoxanthin (Ddx), diatoxanthin (Dtx), and β-carotene (β-Car) contents were quantified by HPLC and expressed as molar ratios to chlorophyll *a* (Chl *a*) (mol/mol). de-epoxidation state of the diadinoxanthin cycle is shown as Dtx/ (Ddx + Dtx), calculated on a molar basis. Values are means ± standard deviation from biological replicates (n = 3).

| Sample (WL) | Fx / Chl *a* | Ddx / Chl *a* | Dtx / Chl *a* | Dtx / (Ddx + Dtx) | β-Car / Chl *a* |
|---|---|---|---|---|---|
| WT | 0.75 ± 0.0061 | 0.33 ± 0.0068 | 0.022 ± 0.00065 | 0.0632 ± 0.00020 | 0.043 ± 0.00010 |
| *Lhcf9-OX 11* | 0.71 ± 0.0035 | 0.31 ± 0.028 | 0.019 ± 0.0026 | 0.0564 ± 0.0022 | 0.043 ± 0.00047 |
| *Lhcf9-OX* 12 | 0.70 ± 0.013 | 0.31 ± 0.024 | 0.021 ± 0.0025 | 0.0604 ± 0.0023 | 0.042 ± 0.00099 |
